# Murine lupus is Neutrophil Elastase-independent in the MRL.Fas*^lpr^* model

**DOI:** 10.1101/857243

**Authors:** Rachael A. Gordon, Jeremy S. Tilstra, Anthony Marinov, Kevin M. Nickerson, Sheldon I. Bastacky, Mark J. Shlomchik

**Affiliations:** Department of Immunology, University of Pittsburgh School of Medicine, Pittsburgh PA; Department of Medicine, University of Pittsburgh School of Medicine, Pittsburgh PA; Department of Pathology, University of Pittsburgh School of Medicine, Pittsburgh PA

## Abstract

Loss of tolerance to nuclear antigens and multisystem tissue destruction is a hallmark of systemic lupus erythematosus (SLE). Although the source of autoantigen in lupus remains elusive, a compelling hypothetical source is dead cell debris that drives autoimmune activation. Prior reports suggest that neutrophil extracellular traps (NETs) and their associated death pathway, NETosis, are sources of autoantigen in SLE. However, others and we have shown that inhibition of NETs by targeting the NADPH oxidase complex and peptidylarginine deiminase 4 (PADI4) did not ameliorate disease in spontaneous murine models of SLE. Furthermore, myeloperoxidase and PADI4 deletion did not inhibit induced lupus. Since NET formation may occur independently of any one mediator, to address this controversy, we genetically deleted an additional important mediator of NETs and neutrophil effector function, neutrophil elastase (ELANE), in the MRL.Fas*^lpr^* model of SLE. ELANE deficiency, and by extension ELANE-dependent NETs, had no effect on SLE nephritis, dermatitis, anti-self response, or immune composition in MRL.Fas*^lpr^* mice. Taken together with prior data from our group and others, these data further challenge the paradigm that NETs and neutrophils are pathogenic in SLE.

## Introduction

SLE is a systemic autoimmune disease characterized by the formation of autoantibodies to nucleic acids and the proteins to which these nucleic acids associate [1]. Loss of tolerance to self-antigens results in immune activation and tissue destruction [1]. Although the origin of autoantigens in SLE are not known, the liberation of antigenic contents from dying cells is considered a likely culprit.

Neutrophils are postulated to play a critical role in SLE pathogenesis by secreting pro-inflammatory cytokines, directly mediating end organ injury, and by forming neutrophil extracellular traps (NETs) [2]. NETs are extruded DNA structures coated with granular and cytoplasmic contents that are released into the extracellular environment. There is substantial disagreement and controversy about the definition of a NET, how to detect and quantify NETs, and what the triggers of and molecular pathways resulting in NET formation are, as summarized in a recent consensus document [3]. These outstanding issues in the NET field make it difficult to study the causative role for NETs in biological processes and diseases.

Classical NET generation in humans and mice relies on NADPH oxidase-generated reactive oxygen species (ROS) [4–6]. However, rapid NADPH oxidase-independent NET formation, nuclear DNA externalization without concomitant cell lysis, and extrusion of mitochondrial DNA have been described [3]. In addition to NADPH oxidase, peptidylarginine deiminase 4 (PADI4) [7–13], neutrophil elastase (ELANE) [10, 14–17], and myeloperoxidase (MPO) [18, 19] have been identified as critical mediators of NET formation.

It is a compelling paradigm that NETs could be a source of autoantigen and a downstream mediator of end-organ damage in SLE. NETs are present in the peripheral blood, skin, and kidneys of SLE patients and mice [2]. Early studies suggested that pharmacological inhibition of PADI4 via pan-PAD inhibition with CL- and BB-CL-Amidine mildly improved clinical manifestations of SLE in murine models [20, 21]. However, the NET hypothesis has recently been challenged by studies that have either genetically deleted or pharmacologically inhibited important NET mediators in multiple murine models of SLE. Genetic deletion of critical NADPH oxidase complex components, required for the neutrophil oxidative burst in addition to ROS-dependent NET formation, exacerbated SLE in mice [22, 23], an observation that also extends to humans [24, 25]. Genetic deletion of *padi4* did not improve clinical or immunological manifestations of SLE in the MRL.Fas*^lpr^* and pristane induced lupus (PIL) mouse models [23, 26]. In fact, disease was exacerbated in the latter [23, 26]. Pharmacological inhibition of the PADI family of enzymes by Cl-amidine had no impact on two inducible models of nephritis [26]. MPO-deficient mice subjected to PIL have increased proteinuria and glomerulonephritis [27]. While these data argue against a role for neutrophils and NETs in SLE pathogenesis, it remains possible that PADI4, CYBB, and MPO independent NETs or other neutrophil effector functions could drive disease.

To address this controversy, it is necessary to use additional genetic approaches to block NET formation. We reason that while the case for NETs driving lupus could posit that one or even more canonical members of the NET cascade would be dispensable specifically in the case of lupus, it would be unlikely that multiple such molecules would all be dispensable. Therefore, to further probe the hypothesis that NETs drive lupus, we decided to genetically target the serine protease *elane,* since multiple studies both directly [10, 14–17] and indirectly [28–31] implicate ELANE in NET formation in mice and humans. ELANE, the second most abundant protein in NETs, is critical for NET formation, functioning by modulating F-actin dynamics and degrading histones important for chromatin decondensation [15]. In addition to its active role in NET formation, ELANE decorates NET structures and is proteolytically active in this setting [32]. Moreover, ELANE is important for neutrophil effector function and can directly mediate tissue damage upon neutrophil degranulation [33]. Therefore, ELANE has the potential to be an effective therapeutic target in SLE.

To directly compare *elane* to both *cybb*- and *padi4*-deficiency in the context of SLE pathogenesis, we elected to eliminate *elane* in the MRL.Fas*^lpr^* mouse model of lupus. The MRL.Fas*^lpr^* model is a leading system to study SLE as it recapitulates nearly all features of the human disease and has accurately predicted responses in human translational studies [34].

Here, we show that genetic deletion of *elane* did not have any impact on clinical or immunological parameters of SLE in MRL.Fas*^lpr^* mice. Taken together with earlier work [22, 23, 26], these findings add additional evidence that challenges the concept that neutrophils and NETs, to the degree that NET generation and neutrophil effector function relies on PADI4 [7–13], ELANE [10, 14–17], or CYBB [4–6], critically drives lupus.

## Materials and Methods

### Mice

*Elane-Cre* C57BL/6 mice were generously provided by Jürgen Roes (Department of Medicine, University College London, London, United Kingdom). Fully backcrossed mice were obtained by crossing *elane^Cre/wt^* allele onto the MRL.Fas*^lpr^* (The Jackson Laboratory catalogue # 000485) background for at least 9 generations. Heterozygous mice were then intercrossed to produce an experimental cohort. SLE pathology was analyzed at 17 weeks for backcrossed cohorts. Mice were genotyped for the *Elane-Cre* allele using a triplex PCR with the following three primers: ELANE-F 5’-AGGGGCACACATCTCTCCAT-3’, ELANE-R 5’-GCCTTCACAGTAACCACCCT, Cre-R 5’-CGCATAACCAGTGAAACAGC-3’. All mice were housed under specific-pathogen-free (SPF) conditions. Animal studies were approved by the University of Pittsburgh Institutional Animal Care and Use Committee.

### Evaluation of SLE pathology

MRL.Fas*^lpr^* SLE cohorts were analyzed as previously described [22, 35]. Skin disease was scored based on the extent of dermatitis on the dorsum of the neck and back. Macroscopic surface area was scored from 0 to 5 for an affected area up to 9.1 cm^2^, with up to 1 additional point for the presence of ear (1/4 point each) and muzzle (1/2 point) dermatitis, as described [36].

Proteinuria was screened using Albustix (Siemens). Plasma was obtained by cardiac puncture. Kidneys were removed, bisected, formalin-fixed, paraffin embedded, and H&E stained. A clinical pathologist (S.I.B.) scored glomerulonephritis on a scale of 1-6 and interstitial nephritis on a scale of 1-4 in a blinded manner [22].

### Flow Cytometry

Flow cytometry was performed as previously described [22]. In brief, spleens and bone marrow were homogenized and red blood cells were lysed using Ammonium-Chloride-Potassium buffer (prepared in house). Cells were resuspended in Phosphate Buffered Saline (PBS) with 3% calf serum and the FcR-blocking antibody 2.4G2. Live/dead discrimination was performed using fixable viability stain 510 (BD) or ethidium monoazide bromide (Invitrogen). Surface and intracellular staining antibodies are listed below. Cells were fixed in 1% paraformaldehyde or Cytofix/Cytoperm (BD) where appropriate. Data were obtained using a LSRII or Fortessa (BD) with FACS DIVA software and analyzed using FlowJo.

### Antibodies used for FACS staining

Antibodies used for FACS surface and intracellular staining were as follows: IA/E-PE (Biolegend, M5/114.15.2), Bst-2-biotin (in-house conjugated, 927), CD11c-PE/Cy7 (BD Pharmigen, HL3), CD45R-APC/Cy7 (BD Pharmigen, RA3-6B2), SiglecH-Al647 (eBioscience, eBio440c), CD19-Pacblue (in-house conjugated, 1D3.2), Ly6G-Al488 (in-house conjugated, 1A8), Gr1-PE/Cy7 (Biolegend, RB6-8C5), Gr1-PE (Biolegend, RB6-8C5), CD11b-APC/Cy7 (Biolegend, M1/70), CD11b-PE (Biolegend, M1/70), F4/80-Al647 (in-house conjugated, BM8), F4/80-APC (Biolegend, BM8), CD44-Al488 (in-house conjugated, 1M7), CD44-APC-Cy7 (Biolegend, 1M7), TcRβ-APC/Cy7, (Biolegend, H57-597), TCRβ-PE/Cy7, (Biolegend, H57-597), CD62L-PE/Cy7 (Biolegend, Mel-14), CD8-Al647 (in-house conjugated, TIB 105), CD4-PE (in-house conjugated, GK1.5), CD138-PE (BD Pharmigen, 281-2), CD19-AI647 (in-house conjugated, 1D3.2), kappa-AI488 (in-house conjugated, 187.1), and Ly6B.2-Fitc (AbD Serotec, 7/4).

### ELISpot assays

AFC producing κ light chain antibodies, IgG1, IgG2a, or IgM were detected by ELISpot as previously described [37]. In brief, 96-well Immulon 4 HBX plates were coated overnight at 4°C with 5 mg/ml polyclonal goat-anti mouse κ (Southern Biotech; 1050-01). Nonspecific binding was blocked with 1% bovine serum albumin in PBS and samples were incubated at 37°C. Alkaline phosphatase-conjugated secondary antibodies (Southern Biotech; Igκ [1050-04], IgG1 [1070-04], IgG2a [1080-04], or IgM [1020-04]) were detected with bromo-4-chloro-3-indolyl phosphate substrate (Southern Biotech).

### ELISAs

Anti-Sm, anti-nucleosome, anti-RNA, rheumatoid factor, total IgM, and IgG ELISAs were performed as previously described [22, 38–41]. Specific antibodies were detected with alkaline phosphatase-conjugated goat anti-mouse IgG (Southern Biotech [1030-04]). The monoclonal antibodies Y2, BWR4, 400tμ23, PL4-2 or PL2-3 (in-house) were used as standards for the anti-Sm, anti-RNA, rheumatoid factor, and anti-nucleosome measurements respectively.

### Antinuclear Antibody Assay (ANA)

HEp-2 immunofluorescence assays were performed as previously described [22]. In brief, serum was diluted 1/200 and then applied to HEp-2 slides (Antibodies Incorporated or BD). Staining was detected using goat anti-mouse IgG FITC at 1:500 (Southern Biotech) ANAs were scored on intensity of nuclear and cytoplasmic staining on a scale of 0–3 and for the presence or absence of mitotic chromatin using wide-field fluorescence microscopy (Olympus IX83; Cellsense software). The scorer used blinded to the genotype and gender of the mice.

### Statistics

Statistical analysis was performed using Prism 7.0 (Graphpad). A two-tailed Mann-Whitney U test, two-tailed Welch’s t test, and Fisher’s Exact test were performed where indicated and appropriate. A p value <0.05 was considered statistically significant.

## Results

To assess the role of *elane*-deficiency in SLE, we utilized the *elane-Cre* allele, which does not produce functional ELANE protein. Homozygous *elane* deficiency does not substantially alter granulocyte development and recruitment [42]. We backcrossed the *elane*-*Cre* allele onto the MRL.Fas*^lpr^* background for 9 generations. Heterozygous mice were then intercrossed to produce an experimental cohort. SLE pathology was analyzed at 17 weeks of age.

### Elane deficiency had no effect on nephritis or dermatitis

No differences in urine protein were detected across *elane* genotypes in either male or female cohorts (Fig 1A). *Elane* deficiency resulted in no significant alterations in glomerulonephritis or interstitial nephritis (Fig 1B). No statistically significant differences in dermatitis were detected among the different *elane* genotypes in either cohort (Fig 1C). Similarly, *elane* genotype did not alter splenomegaly or lymphadenopathy (Fig 1D).

**Fig 1.** *Elane*-genotype does not impact lupus nephritis, dermatitis, or lymphadenopathy/splenomegaly. **(A)** Proteinuria scores. **(B)** Glomerulonephritis (left panel) and Interstitial nephritis (right panel) scores (*elane^+/+^* females n=17). **(C)** Dermatitis scores. **(D)** Axillary lymph node (left panel) and spleen weights (right panel). Scores and weights are represented as a function of *elane-*genotype and gender at 17 weeks of age. Bars represent the median ± interquartile range. A two-tailed Mann-Whitney U test was performed to determine statistical significance within each gender (*elane^-/-^* males n=9; *elane^+/+^* males n= 13; *elane ^-/-^* females n=20; *elane ^+/+^* females n=18 mice per group unless otherwise indicated).

### Elane genotype has no impact on loss of tolerance and the anti-self response

To determine the effect of ELANE on the autoantibody response in SLE, we characterized the ANA patterns in *elane*-sufficient and -deficient MRL.Fas*^lpr^* mice. No differences were identified in the dominant ANA pattern or staining intensity across groups (Supplemental Fig 1A). ELANE deficiency did not alter anti-chromatin autoantibodies as no differences in mitotic chromatin staining were observed (Supplemental Fig 1B). Concordant with these data, we did not detect any changes in anti-RNA antibody (Fig 2A), anti-Sm antibody (Fig 2B), anti-nucleosome antibody (Fig 2C), and rheumatoid factor (Fig 2D) titers by ELISA among the groups.

**Fig 2.** *Elane*-genotype does not significantly alter the anti-self response or the AFC compartment. **(A-D)** Serum anti-RNA (A), anti-Sm (B), anti-nucleosome (*elane^+/+^* males n= 11; *elane ^+/+^* females n=16 mice per group) (C), and rheumatoid factor (D) antibody titers at 17 weeks of age. **(E)** Numbers of Igκ^+^ antibody forming cells (AFCs) per spleen as determined by ELISpot (left panel). Percentages of live cells that are TCRβ^-^ CD44^+^ CD138^+^ intracellular κ^+^ AFCs in spleens (right panel). **(F-H)** Numbers of IgG1 (F), IgG2a (G), and IgM (H) AFCs per spleen as determined by ELISpot (*elane^-/-^* males n=8; *elane^+/+^* males n= 11; *elane ^-/-^* females n=19; *elane ^+/+^* females n=13 mice per group unless otherwise indicated). Data representation and statistics are as in Fig 1 unless otherwise indicated. In panel E (right), bar graphs are represented as the mean ± SEM and a two-tailed Welch’s T test was performed to determine statistical significance within each gender.

*Elane*-genotype did not affect the percentages of CD19^+^ B cells (Fig 3D) or CD19^low-int^ CD44^+^ CD138^+^ intracellular k^high^ AFCs (Fig 2E, right panel). Similarly, there were no statistically significant differences in total kappa Ig AFC ELISpots (Fig 2E, left panel). Wild-type female mice had a higher number of IgG1 AFCs compared to their *elane*-deficient MRL.Fas*^lpr^* counterparts (Fig 2F, p=0.0141). This finding is unlikely to be biologically significant, as these data were not mirrored across gender. No differences were identified in IgG2a and IgM AFC ELISpots amongst the groups (Fig 2G & H).

**Fig 3.** *Elane*-genotype does not substantially affect the myeloid, DC, or T cell compartments. **(A&B)** Percentages of live CD11b^+^ Ly6G^+^ neutrophils (left panel) and CD11b^+^ GR1^low-int^ F4/80^+^ macrophages (right panel) in the bone marrow (A) and spleens (B). **(C)** Percentages of live CD19^-^ MHCII^+^ CD11c^+^ conventional dendritic cells (DCs) (left panel) and CD19^-^ BST2^+^ CD11c^+^ plasmacytoid DCs (right panel). **(D)** Percentages of live CD19^+^ total B cells (left panel) and TCRβ^+^ total T cells (right panel). **(E)** Percentages of live TCRβ^+^ CD4^+^ T cells (left panel) and of CD4^+^ CD44^+^ CD62L^-^ activated T cells (right panel). **(F)** Percentages of live TCRβ^+^ CD8^+^ T cells (left panel) and of CD8^+^ CD44^+^ CD62L^-^ activated T cells (right panel). Bar graphs represent the mean and error bars ± SEM. A two-tailed Welch’s t-test was performed to determine statistical significance within each gender. (*elane^-/-^* males n=9; *elane^+/+^* males n= 13; *elane ^-/-^* females n=20; *elane ^+/+^* females n=18 mice per group unless otherwise indicated).

### ELANE deficiency did not substantially change the myeloid compartment

*Elane* genotype had only a minor impact on the myeloid compartment in MRL.Fas*^lpr^* mice. The percentages of CD11b^+^Ly6G^+^ (Fig 3A, left panel) bone marrow (BM) neutrophils and CD11b^+^F4/80^+^Gr1^low-int^ BM macrophages (Fig 3B, right panel) were not statistically different among the groups. Additionally, no changes in CD11b^+^Ly6G^+^ (Fig 3B, left panel) splenic neutrophils or CD11b^+^F4/80^+^Gr1^low-int^ (Fig 3B, right panel) splenic macrophages were identified across *elane* genotypes. Female wild-type mice had a greater percentage of splenic cDCs compared to the *elane^-/-^* group (Fig 3C, left panel, p=0.0039). No differences in the percentages of CD19^-^BST2^+^ SiglecH^+^ pDCs (Fig 3C, right panel) were identified among *elane* genotypes.

### ELANE deficiency had a modest impact on the lymphoid compartment

*Elane* deficiency did not substantially impact the lymphoid compartment. All genotypes exhibited indistinguishable total percentages of TCRβ^+^ T cells (Fig 3D). The percentages of CD4^+^ T cells were elevated in female knock-out mice (Fig 3E, left panel, p=0.0429) while CD4^+^CD44^+^CD62L^-^ activated T cells were elevated in male wild-type mice (Fig 3E, right panel, p=0.0348). The percentages of CD8^+^ T cells and CD8^+^CD44^+^CD62L^-^ activated T cells were similar amongst wild-type and *elane*-deficient mice (Fig 3F).

## Discussion

The data from the genetic study presented here in a relevant murine model of lupus does not support the hypothesis that ELANE, and by extension ELANE-dependent NET generation and neutrophil effector function, alters immune system composition or pathology in SLE. This conclusion is reinforced by prior work from our group and that of others targeting PADI4 in SLE and nephritis models that yielded similar results [23, 26]. Moreover, NADPH deficiency exacerbates SLE in mice and humans. It remains unknown whether this phenotype is neutrophil or NET dependent as NADPH oxidase is implicated in immunoregulatory functions in both the myeloid and lymphoid compartments [43].

In humans, *elane* mutations are generally associated with congenital neutropenias but not systemic autoimmunity [44]. However, a recent study identified a patient with a novel loss of function polymorphism (G210R) in exon 5 of *elane* that did not confer congenital neutropenia, but the patient did exhibit inflammatory arthritis and recurrent parvovirus infections [14]. Concordant with previous observations, NET formation was defective in this patient. Moreover, patients with Papillon-Lefèvre syndrome (PLS), characterized by mutations that inactivate Cathepsin C, a cysteine protease that processes ELANE into its active state [30, 45], develop severe periodontitis. While systemic autoimmune syndromes are not typically associated with PLS, a pediatric patient with PLS who subsequently developed SLE has been reported [46]. Similarly, NET generation was found to be defective in PLS patients [30, 45]. Loss of function mutations in endogenous ELANE inhibitors, such as secretory leukocyte protease inhibitor and Serpin Family B Member 1, are associated with increased NET formation but their role in systemic autoimmunity has not been extensively investigated [28, 29].

It was conceivable that ELANE protease activity could cleave self-antigens, thus impacting their immunogenicity. Proteases have been suggested as sources to generate novel immunogenic forms of self-antigens [47]. However, ELANE deficiency did not alter the anti-self response nor clinical parameters of SLE in the MRL.Fas^l*pr*^ model, ruling out one possible source of proteases that could fuel such a mechanism of loss of tolerance in autoimmunity.

NADPH oxidase-deficient mice subjected to a monosodium urate crystal induced model of chronic neutrophilic inflammation, a model of gout, developed more severe arthritis [48]. This phenotype was attributed to the inability of NADPH oxidase-deficient mice to generate aggregated NETs to degrade proinflammatory mediators. Moreover, the adoptive transfer of aggregated NETs into these animals reduced disease severity [48]. Aggregated NETs can degrade cytokines in an ELANE-dependent manner, as treatment of these structures with multiple elastase inhibitors prevented the degradation of cytokines and chemokines [48]. These data coupled with the exacerbated disease observed in *cybb*-deficient SLE prone mice and *padi4*-deficient mice subjected to the PIL model suggests a homeostatic function for NETs in autoimmunity.

While many groups have implicated ELANE as a critical mediator of NET formation [10, 14–17, 28–31], a few reports suggest that NETs can occur in the absence of ELANE. In one report, immune complexes induced ELANE-independent and FcgrIIA-dependent NET formation in normal neutrophils [49]. Another report concluded that neutrophils from *elane*-deficient mice or wild-type neutrophils pretreated with ELANE inhibitors did form NETs in response to canonical NET inducing stimuli, such as PMA and calcium ionophores [50]. In this study, *elane-*deficient mice formed NETs in an experimental model of deep venous thrombosis (DVT) and ELANE deficiency did not protect mice from DVT in this model [50]. Since PADI4 deficiency and pharmacological inhibition reduces NET formation in lupus and PADI4 pharmacological inhibition or genetic deletion does not impact SLE pathogenesis [23, 26], and given the large number of reports implicating ELANE in NET formation, it seems unlikely that ELANE-independent NETs drive autoimmunity in this setting.

The multiple studies that blocked or deleted mediators of NET formation but which did not observe ameliorated disease raise a broader question as to the role of neutrophils per se in lupus pathogenesis. Infiltration of neutrophils into the kidney has been identified in both SLE patients and in multiple murine models of the disease [51]. Despite these associations, it is striking that nearly 20-50% of SLE patients are neutropenic [52, 53] and/or have abnormal neutrophil function. SLE neutrophils display reduced superoxide production [54] and phagocytosis [55]; however, low density granulocytes (LDGs)—a neutrophil subset enriched in SLE patients—produce type I interferons and proinflammatory cytokines [56]. Animal models of lupus have not supported a pathogenic role for neutrophils in SLE pathogenesis; rather, data has supported a regulatory or neutral role for neutrophils. Antibody mediated depletion of neutrophils early in disease *increased* autoantibody titers and glomerulonephritis in the NZB/W model [57, 58]. However, neutrophil depletion did not alter autoantibody responses nor renal pathology in established disease [57]. Conditional deletion of neutrophils in mice subjected to the PIL model resulted in increased antinuclear antibody responses [23].

The major message from the current studies comes from interpreting them in the context of other genetic and inhibitor studies that have probed whether NETs and neutrophils promote lupus. Our work adds ELANE to the list of CYBB and PAD4 as proteins that are required for NET formation, yet are not needed for lupus. While in each case, it could be argued that the type of NET generation seen in SLE proceeds independently of the protein in question, it seems unlikely that the putative type of NET generation that is hypothesized to critically drive lupus is independent of three of the major pathways documented to be important for NET formation. This point, which is punctuated by the current data, along with the lack of a role for neutrophils per se in driving lupus, heralds a juncture at which the field should seriously reevaluate whether the “NETs drive lupus” hypothesis remains valid and whether blocking NET formation or neutrophil function makes sense as a therapeutic strategy. In contrast, multiple studies have shown regulatory roles in various systems for CYBB, and to an extent PAD4 and MPO, such that disease is exacerbated in their absence [22, 23, 26, 27, 59]. We would thus posit that neutrophils and their effector mechanisms, such as NET generation, may have evolved to protect us from autoimmune diseases, rather than function as vectors to promote them.

## Acknowledgements

This study utilized the NIH-sponsored Pittsburgh Center for Kidney Research.

## Supporting Information

**S1 Fig. Supplemental Figure 1.**
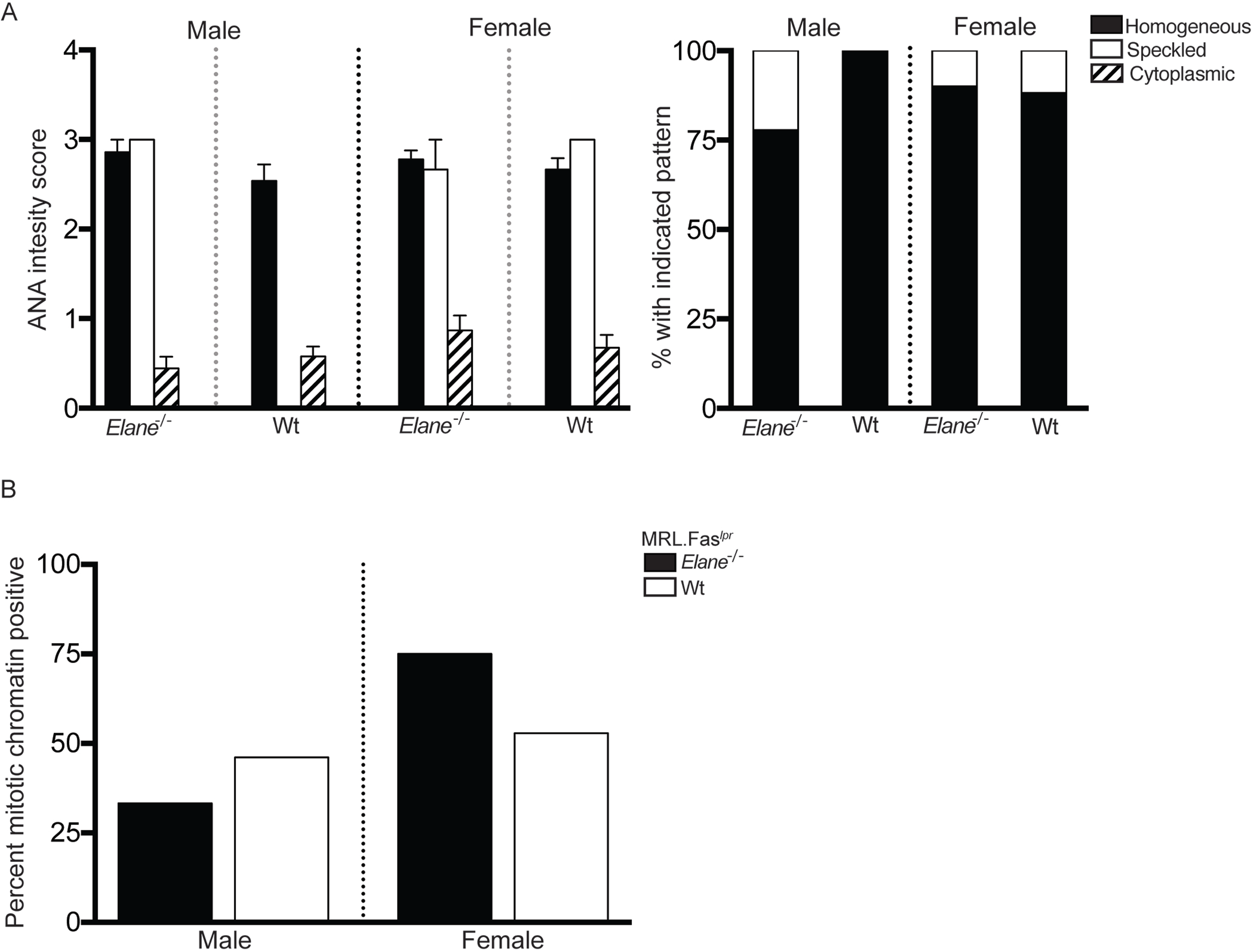
*Elane*-genotype does not alter ANA patterns. (A) HEp2-ANA slides were scored for intensity of nuclear and cytoplasmic staining patterns (left panel). Data representation and statistics are as in figure 2F unless otherwise indicated. Dominant ANA pattern classified as nuclear (homogenous), nuclear (speckled), or cytoplasmic (right panel). (B) HEp2-ANA slides were scored for the presence or absence of mitotic chromatin staining. In panel B, a Fisher Exact test was performed to determine statistical significance within each gender (*elane^-/-^* males n=9; *elane^+/+^* males n= 13; *elane ^-/-^* females n=20; *elane^+/+^* females n=17 mice per group).

